# The integrated stress response promotes macrophage inflammation and migration in autoimmune diabetes

**DOI:** 10.1101/2024.10.31.621343

**Authors:** Jiayi E. Wang, Charanya Muralidharan, Titli Nargis, Jacob R. Enriquez, Ryan M. Anderson, Raghavendra G. Mirmira, Sarah A. Tersey

## Abstract

Type 1 Diabetes (T1D) is an autoimmune disease characterized by the destruction of insulin producing pancreatic β-cells. Macrophages infiltrate islets early in T1D pathogenesis, preceding the influx of T- and B-lymphocytes. The integrated stress response (ISR), a cellular pathway activated during stress, coordinates adaptive changes in gene expression to maintain cell function and survival. To study the ISR in macrophages, bone-marrow derived macrophages were treated with a pharmacological inhibitor of the ISR (ISRIB) and polarized to a proinflammatory M1-like state. We found a reduction in the number of proinflammatory macrophages as well as a reduction in iNOS mRNA and protein levels with ISRIB treatment. RNA-sequencing revealed a reduction in pathways related to stress responses, including ER stress, reactive oxygen species (ROS) regulation, and autophagy as well as migration pathway genes. ISRIB treatment led to decreased macrophage migration after stimulation in vitro and decreased macrophages to the site of injury after tailfin cut in zebrafish in vivo. Pre-diabetic female non-obese diabetic (NOD) mice administered ISRIB demonstrated an overall reduction in the macrophage numbers in the pancreatic islets. Additionally, the insulitic area of pancreata from ISRIB treated NOD mice had increased PD-L1 levels. Our study provides new insights into ISR signaling in macrophages and demonstrates that the ISR might serve as a potential target for intervention in macrophages during early T1D pathogenesis.

## INTRODUCTION

Type 1 diabetes (T1D) is an autoimmune disease characterized by insulitis, an inflammatory process targeting the insulin-producing pancreatic β cells. This destruction leads to diminished insulin production and hyperglycemia (1). During the course of T1D, innate immune cells, particularly macrophages, are the first to infiltrate the pancreatic islets (2). Macrophages adopt distinct activation states (M1/M2) influenced by tissue microenvironments. The proinflammatory M1 macrophages contribute to the early stages of insulitis and establish a critical link between stressed β cells and the activation of the adaptive immune system. Depleting macrophages in non-obese diabetic (NOD) mice prevents insulitis and the subsequent development of diabetes (3, 4). Similarly, in a zebrafish model of chemically induced β-cell injury, macrophage depletion reduced β-cell loss and mitigated hyperglycemia (5). However, systemic depletion of macrophages is impractical due to its detrimental effects on overall immunity. Therefore, strategies aimed at modulating macrophage function could potentially be used to treat or prevent T1D. For example, our previous research demonstrated that deleting *Alox15,* a key proinflammatory enzyme, in macrophages prevented autoimmune diabetes in NOD mice by reducing proinflammatory macrophage migration (5). In T1D, macrophages predominantly exhibit a M1 proinflammatory phenotype, which intensifies inflammation by secreting inflammatory cytokines like IL-1β, presenting antigens via MHC class II molecules to recruit adaptive immune cells, and generating reactive oxygen species (ROS) that exacerbate oxidative stress (2).

The integrated stress response (ISR) is a signaling network triggered in response to environmental and pathological stressors leading to suppression of mRNA translation to restore cellular energy balance. The ISR is mediated by four key kinases: PERK (responding to endoplasmic reticulum stress), PKR (activated by viral infections), HRI (responsive to heme deficiency), and GCN2 (triggered by amino acid deprivation). Activation of these kinases leads to the phosphorylation of the eukaryotic translation initiation factor 2 alpha (eIF2α) at the serine 51 residue, which suppresses global translation initiation as an adaptive response (6). Stressors capable of activating the ISR are linked with T1D development and pathology. In human pancreata from pre-T1D donors with positive autoantibodies, 20 ISR-associated genes are significantly altered, including the genes encoding three of the four ISR kinases - PERK, PKR, and GCN2 (7). Indeed, inhibiting the ISR in human islets has been shown to confer protection against harmful effects of proinflammatory cytokines (8, 9). Notably, inhibition of the ER-induced PERK kinase, a shared arm of both the unfolded protein response (UPR) and ISR, has been shown to reduce islet inflammation and delay autoimmune diabetes in NOD mice. In addition, pharmacological inhibition of ISR through a previously described inhibitor, ISRIB, in NOD mice resulted in significant reduction in insulitis (8). Despite these findings, the specific functional effects of the ISR on macrophages during the chronic inflammation of T1D remain unclear. In this study, we utilized ISRIB to delineate the effects of ISR in macrophage function using in vitro macrophages, zebrafish, and NOD mice. Our findings provide evidence that ISR promotes macrophage migration and proinflammatory phenotype in the context of T1D.

## RESULTS

### ISR regulation of macrophage mRNA translation in T1D pathogenesis

Macrophages play a critical role in autoimmune diabetes development. In the NOD mouse model of T1D, about 80% of the females develop spontaneous hyperglycemia by 18 weeks of age (10). To understand the temporal dynamics of the macrophage population in the pathogenesis of T1D, pancreata of pre-diabetic female NOD mice were immunostained for the macrophage marker F4/80 (Figure 1A). We observed a significant increase in macrophage numbers from 6 to 8 weeks of age, which remained elevated through 12 weeks of age (Figure 1B). These increases coincide with the period before the onset of diabetes in female NOD mice, typically occurring between 12 and 20 weeks (10). During this pre-diabetic period, immune infiltration into the pancreatic islets primes the environment for T1D onset. The marked expansion of macrophage populations, particularly from 6 to 8 weeks, confirms their involvement in the early inflammatory processes associated with T1D development, suggesting a critical window for therapeutic intervention targeting macrophages.

**Figure 1:**
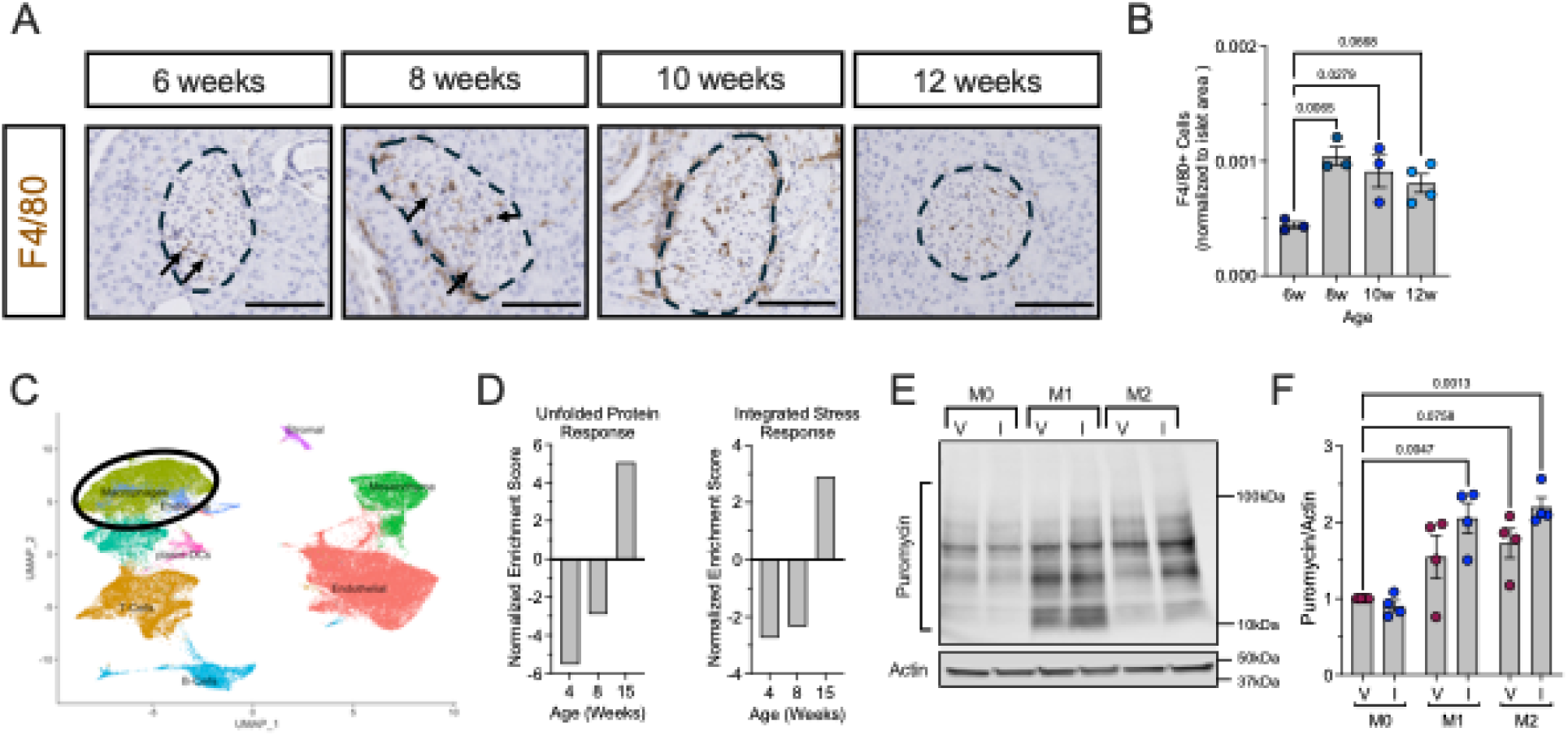
ISR Regulates Macrophage Translation in T1D Pathogenesis. (**A**) Representative images of pancreata from female NOD mice overtime stained for F4/80 (macrophage marker; brown) and counterstained with hematoxylin (blue). *Dotted lines* indicate islets and *arrows* indicate macrophages. Scale bars: 100 μm. (**B**) Quantification of F4/80+ cells in islet area; *n* = 3-4 biological replicates. (**C**) UMAP embeddings of a reanalysis of scRNA-Seq of islets from female NOD mice. (**D**) GSEA of the macrophage cell population identified in **C** for UPR (*left panel*) and ISR (*right panel*). (**E**) Representative immunoblot of puromycin incorporation normalized to Actin in vehicle or 50 nM ISRIB treated unpolarized (M0) and polarized (M1 or M2) BMDMs isolated from 8-week-old male *C57B/6J* mice; *n* = 4 biological replicates. (**F**) Quantification of the immunoblots in **E**. Statistical tests: ANOVA. Data are represented as mean ± SEM.

Given that the unfolded protein response (UPR) and ER stress in β cells is a well-established factor in T1D pathogenesis (10, 11), we sought to demonstrate that the UPR and ER stress also plays a role in macrophages. To do this, we reanalyzed a publicly available single-cell RNA sequencing dataset from islets of female pre-diabetic NOD mice at 4, 8, and 15 weeks of age (12), stratifying the macrophage population (Figure 1C). Gene Set Enrichment Analysis (GSEA) revealed a gradual enrichment of genes of the UPR with advancing age, indicating increased ER stress and accumulation of unfolded or misfolded proteins in macrophages (Figure 1D). Additionally, GSEA showed an enrichment of genes involved in the ISR in macrophages over time (Figure 1D). While the UPR is primarily activated to restore normal ER function by managing the load of unfolded proteins, the ISR coordinates a broader cellular response to various stress signals, including ER stress, to regulate protein translation through eIF2α phosphorylation (p-eIF2α).

To assess the impact of the ISR on global protein translation, we employed the surface sensing of translation (SUnSET) technique, which detects newly synthesized proteins via the incorporation of puromycin into elongating polypeptide chains (13). Bone marrow-derived macrophages (BMDMs) were isolated from 8-week-old male *C57BL/6J* mice and treated with either vehicle or 50 nM ISRIB, an ISR inhibitor (14). Macrophages were then polarized overnight into three states: a proinflammatory M1-like state using LPS and IFN-γ, an anti-inflammatory M2-like state using IL-4, or left as unpolarized M0 control macrophages. Puromycin-labeled nascent peptides were visualized through immunoblotting with anti-puromycin antibodies, allowing for quantification of overall protein synthesis (Figure 1E). We observed significantly increased puromycin incorporation in M1- and M2-polarized macrophages compared to M0 controls similar to the data we have previously observed using a polyribosome profiling technique (15). ISR inhibition with ISRIB further enhanced translation in both polarized states, showing an upward trend despite not reaching statistical significance (Figure 1F). These results suggest that ISR inhibition enhances protein translation in both pro- and anti-inflammatory macrophages, which are pivotal in different stages of inflammation during T1D progression.

### ISR inhibition reduces proinflammatory macrophage polarization

To further elucidate the role of the ISR in macrophage function, we next investigated whether the ISR influences macrophage polarization toward proinflammatory (M1-like; LPS and IFN-γ) or anti-inflammatory (M2-like; IL-4) phenotypes using flow cytometry, RNA markers, and protein markers (16). BMDMs were isolated from 8-week-old male *C57BL/6J* mice and treated with either vehicle or 50 nM ISRIB prior to stimulation to M1- or M2-like states. Flow cytometry analysis showed that macrophages from the ISRIB-treated group exhibited reduced propensity to polarize towards the M1-like phenotype, as indicated by decreased expression of the M1 marker inducible nitric oxide synthase (iNOS) (Supplemental Figure S1A and Figure 2A). Additionally, *Nos2* (encoding iNOS) and *Il6* mRNA levels were downregulated, with *Nos2* showing a significant decrease in ISRIB-treated BMDMs (Figure 2B and C). This downregulation was further confirmed at the protein level, as immunoblotting revealed a reduction in iNOS production in ISRIB-treated macrophages (Figure 2D). In contrast, vehicle- and ISRIB-treated BMDMs showed a similar capacity to polarize to the M2-like anti-inflammatory state, as assessed by flow cytometry of the M2 marker CD206 (Supplemental Figure S2B and S2C). These results suggest that ISR inhibition dampens the proinflammatory M1 polarization of macrophages. Since the polarization potential of M2 macrophages remained unaffected by ISRIB-mediated ISR inhibition, the subsequent phases of the study focused on the M1 proinflammatory macrophages, which are relevant to the pathogenesis and onset of T1D.

**Figure 2:**
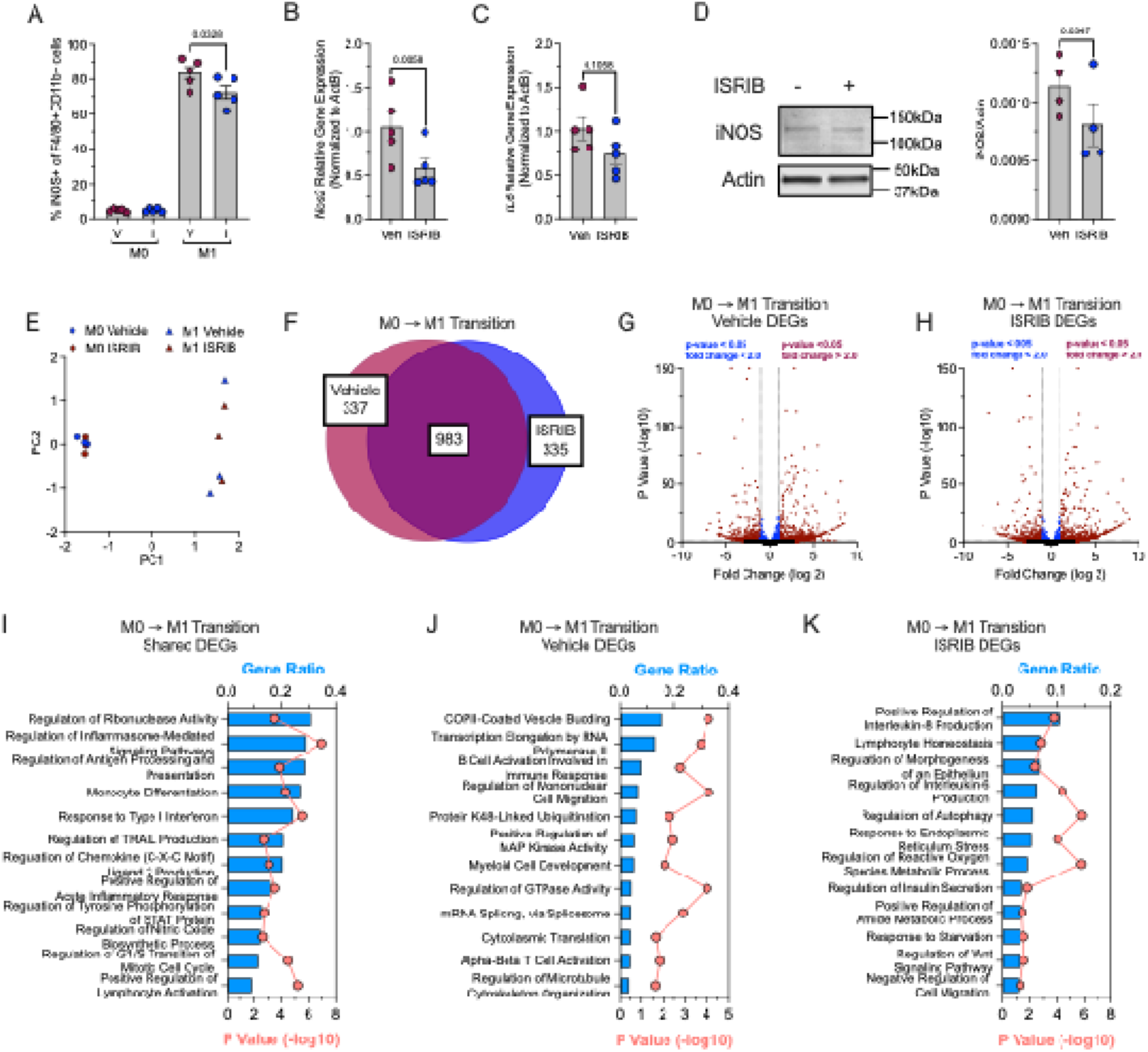
ISR Inhibition Reduces Proinflammatory Macrophages. BMDMs from male 8-week-old *C57Bl/6J* mice were unpolarized (M0) or polarized to M1-proinflammatory macrophages in the presence or absence of 50 nM ISRIB. **(A)** Flow cytometry analysis of iNOS+ (M1-polarization marker) cells as a percentage of F4/80+ CD11b+ cells; *n* = 5 biological replicates. (**B**) Relative *Nos2* mRNA levels measured by qRT-PCR normalized to *Actb* in M1-polarized BMDMs; *n* = 5 biological replicates. (**C**) Relative *Il6* mRNA levels measured by qRT-PCR normalized to *Actb* in M1-polarized BMDMs; *n* = 5 biological replicates. (**D**) Representative immunoblot showing iNOS and Actin in M1-polarized BMDMs (*left panel*) and quantification of iNOS levels; *n* = 4 biological replicates. Statistical tests: Paired t-tests or ANOVA. Data are represented as mean ± SEM. (**E**) Principal component analysis plot of RNA-sequencing results from vehicle- or ISRIB-treated unpolarized or M1-polarized BMDMs; *n* = 3 biological replicates. (**F**) Shared and uniquely expressed genes identified between vehicle- and ISRIB-treated BMDMs during M0 to M1-like transition phase. Venn diagram generated using BioVenn. (**G**) Volcano plot of differentially expressed genes in M0 to M1-like transition in vehicle-treated group. (**H**) Volcano plot of differentially expressed genes in M0 to M1-like transition in ISRIB-treated group. (**I**) Gene ontology (GO) pathway analysis of differentially expressed genes commonly regulated during M0 to M1-phase in both vehicle and ISRIB treatment groups. (**J**) GO pathway analysis of differentially expressed genes in M0 to M1-phase exclusive to vehicle group. (**K**) GO pathway analysis of differentially expressed genes in M0 to M1-phase exclusive to ISRIB group.

### RNA-sequencing revealed changes in pathways associated with protein translation and cell migration after ISR inhibition

To explore the effects of ISR intervention on macrophage gene expression, bulk RNA sequencing was conducted on BMDMs from *C57BL/6J* mice in the presence or absence of ISRIB. BMDMs were unpolarized (M0) or polarized to a proinflammatory M1-state. Principal component analysis (PCA) of the transcriptomic data revealed distinct clustering of macrophages based on their polarization state (Figure 2E). There were no significant transcriptomic differences between M0 states of the vehicle and ISRIB-treated macrophages, suggesting no effect of ISR inhibition in unpolarized macrophages. Next, we analyzed the gene expression changes involved in the transition from the unpolarized M0 state to the proinflammatory M1-like state in BMDMs treated with either vehicle or ISRIB. The analysis identified 983 differentially expressed genes (fold-change, FC, >2, p <0.05) that were common between both treatment groups (Figure 2F-H). Gene ontology (GO) pathway analysis of these shared genes revealed consistent alterations in terms cytokine and chemokine signaling, cellular differentiation and activation, as well as antigen processing and presentation between vehicle- and ISRIB-treated M1 transitions (Figure 2I). This consistency suggests that core immune and inflammatory processes remain active during M1 polarization and are largely unaffected by ISR inhibition.

337 genes were significantly differentially altered in vehicle-treated macrophages, while 335 genes were altered in ISRIB-treated macrophages. The 337 genes altered with vehicle treatment mapped to GO pathways such as B cell and T cell activation, myeloid cell development, and mononuclear cell migration reflecting the immune system’s mobilization and preparation for inflammatory action (Figure 2J). The terms “transcription elongation by RNA polymerase II” and “cytoplasmic translation” suggest an effect on gene expression and protein synthesis necessary for the transition to a proinflammatory state. Given that the ISR modulates protein translation under stress conditions, these findings underscore the critical role of ISR in M1 proinflammatory macrophages. The 335 genes altered with ISRIB treatment mapped to pathways involved in stress response, including ER stress, reactive oxygen species (ROS) regulation, and autophagy (Figure 2K). Notably, there was also enrichment for negative regulation of cell migration, indicating that ISR inhibition may reduce macrophage mobility.

Further analysis of the NOD single-cell RNA sequencing dataset revealed that genes associated with the positive regulation of macrophage migration were most enriched at 4 weeks, maintained similar levels at 8 weeks, but became significantly downregulated by 15 weeks (Supplemental Figure S2A). This pattern suggests that macrophage migratory capacity is more pronounced during the early stages of T1D development but diminishes as the disease progresses, when macrophages have already infiltrated the islets.

### ISR inhibition impairs migration of proinflammatory macrophages

In light of the negative regulation of cell migration observed in the transcriptomics following ISR inhibition, we evaluated the mRNA expression of a panel of macrophage migratory genes. Among the genes assessed, *Cxcr3*, *Ccr2*, and *Lgals3* were significantly downregulated in ISRIB-treated M1-macrophages, while *Mif* and *Cxcr1* exhibited a trend towards reduced expression (Figure 3A). The expression levels of *Ccr4*, *Ccr7*, *Ccl5*, and *Slamf*9 remained unaffected by ISRIB treatment (Supplemental Figure S2B). We next assessed the impact of ISR inhibition on the migratory capabilities of proinflammatory macrophages in vitro using a transwell migration assay. We quantified the migration of polarized BMDMs by measuring the number of cells that traversed the transwell insert. Polarization to the M1-like state enhanced macrophage migration over M0 macrophages. ISRIB treatment of M1-like macrophages exhibited reduced migratory activity (Figure 3B).

**Figure 3:**
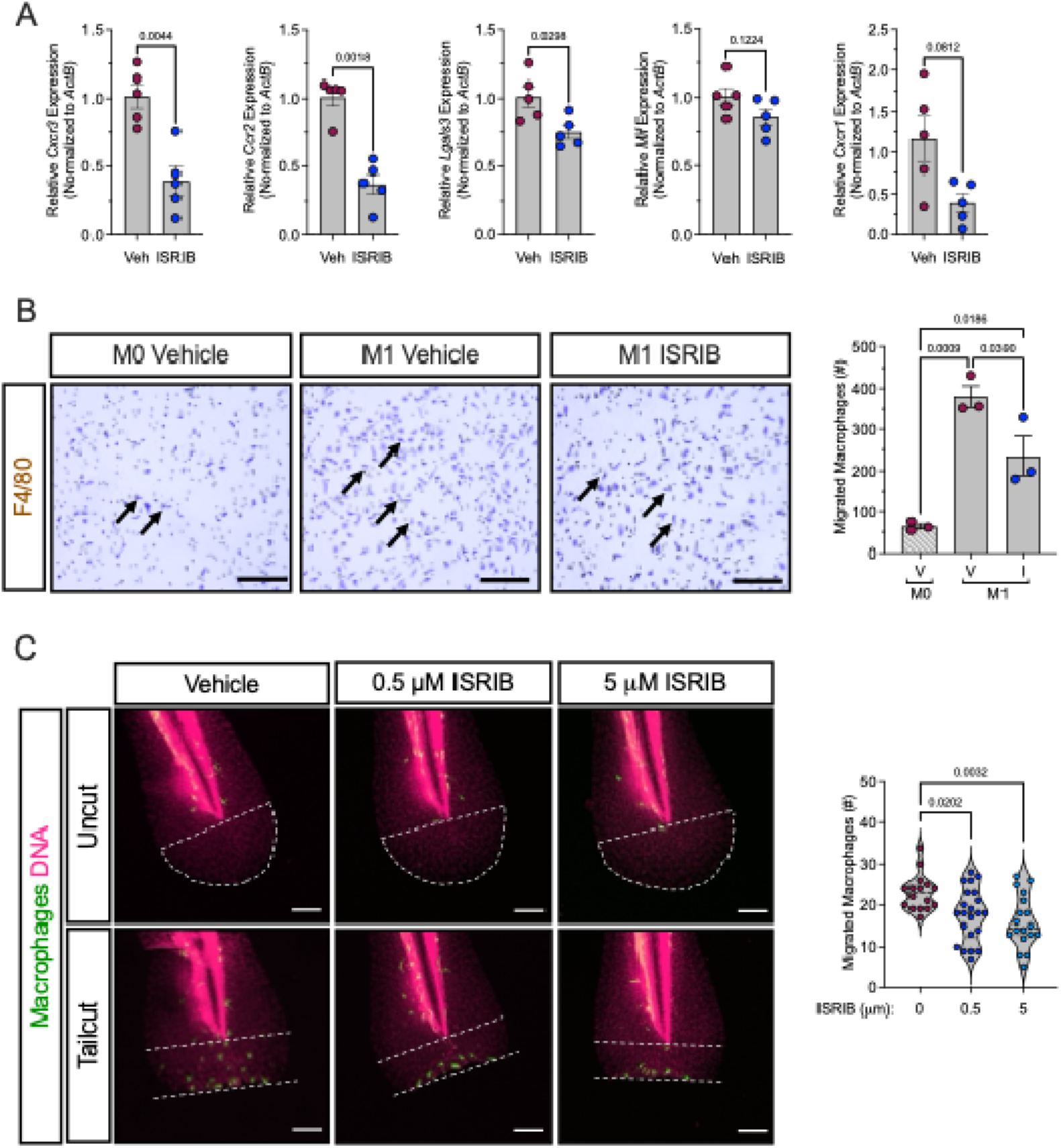
ISR Inhibition Impairs Migration of Proinflammatory Macrophages. BMDMs from male 8-week-old *C57Bl/6J* mice were unpolarized (M0) or polarized to M1-proinflammatory macrophages in the presence or absence of 50 nM ISRIB. (**A**) Relative mRNA levels of key migration markers measured by qRT-PCR normalized to *Actb* in M1-polarized BMDMs; *n* = 5 biological replicates. (**B**) Representative images of in vitro Boyden chamber macrophage migration (stained with crystal violet) of unpolarized (M0) or M1-polarized macrophages in the presence or absence of ISRIB (*left panels)*. Arrows indicate macrophages (violet). Quantification of the number of migrated macrophages (*right panel*); *n* =3 biological replicates. Scale bars:100μm. (**C**) Tg(mpeg:eGFP) zebrafish were treated with vehicle, 0.5 μM or 5 μM ISRIB and then underwent tailfin injury at 3 dpf. Representative images of uncut and cut zebrafish tails showing GFP-labeled macrophages (green) and TO-PRO3-labeled nuclei (magenta) (*left panels*). Quantitation of relative number of migrated macrophages (*right panel)*. Dotted lines indicate the quantified tailfin region for macrophage counts, extending from the area distal to the notochord to the injury site); *n =* 16-22 fish per condition. Statistical tests: Paired t-tests or ANOVA. Data are represented as mean ± SEM.

To explore the effects of ISR inhibition on migration within a more complex inflammatory environment, we utilized an in vivo zebrafish tailfin injury model to assess macrophage migration (5). Zebrafish develop a fully functional innate immune system by 2-3 days post-fertilization (dpf), and owing to their optical transparency and the ease of generating transgenic lines, they serve as an excellent model for studying immune responses in vivo (*17*). We utilized the well-characterized Tg(mpeg) transgenic zebrafish line, in which macrophages are labeled with green fluorescent protein (GFP) (18). Embryos were treated with 0.5 and 5 μM ISRIB or DMSO vehicle at 2 dpf for 18 hours prior to a mechanical tailfin injury. In the absence of injury, macrophage migration was not observed in either vehicle- or ISRIB-treated embryos, indicating that the number of macrophages observed in the tailfin following injury reflects the inflammatory stimulus, rather than baseline activity (Figure 4C). The number of macrophages migrating to the injury site 6h post tailfin cut was reduced in both ISRIB-treated groups compared to vehicle-treated controls (Figure 4C). These findings align with our in vitro data, reinforcing the conclusion that ISR inhibition impairs the migration of proinflammatory macrophages.

**Figure 4:**
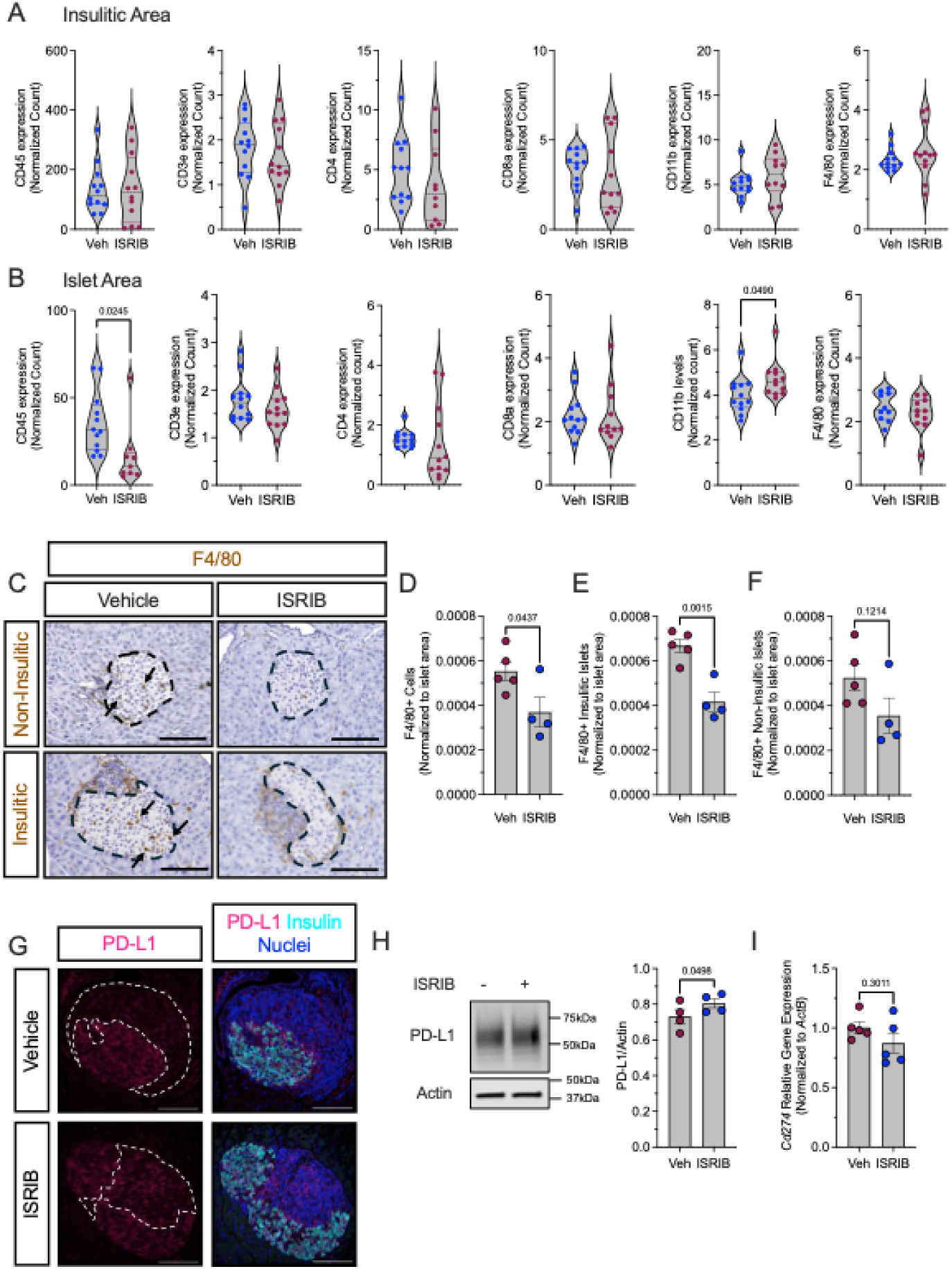
ISR Inhibition Reduces Islet Macrophages and increases PD-L1 levels in female NOD mice. Prediabetic 6-week-old female NOD mice were treated with ISRIB (2.5 mg/kg) for two weeks. (**A**) Identified protein levels in the insulitic area. (**B**) Identified protein levels in the insulin-positive area; *n* = 10-12 ROI from 2 mice per group. (**C**) Representative images of pancreata from ISRIB-treated NOD mice stained for F4/80 (brown), and counterstained with hematoxylin (blue). *Dotted lines* indicate islets and *arrows* indicate macrophages. Scale bars: 100μm. (**D**) Quantification of F4/80+ cells in islet area; *n* = 4-5 biological replicates. (**E**) Quantification of F4/80+ cells in islets with insulitis. (**F**) Quantification of F4/80+ cells in islets without insulitis. (**G**) Representative images of pancreata from vehicle- or ISRIB-treated NOD mice stained for PD-L1 (magenta), insulin (cyan), and nuclei (blue). *Dotted lines* indicate insulitic regions. Scale bars: 100μm. (**H**) Representative immunoblot showing PD-L1 and Actin in M1-polarized BMDMs treated with vehicle or ISRIB (*left panels*) and quantification of PD-L1 levels (*right panel*); *n* = 4 biological replicates. (**I**) Relative *Cd274* mRNA levels measured by qRT-PCR normalized to *Actb* in M1-polarized BMDMs; *n* = 5 biological replicates. Statistical tests: Unpaired t-tests **A-B, D-F** and paired t-tests **H-I**. Data are represented as mean ± SEM.

### ISR inhibition reduces macrophage number in pancreatic islets in NOD mice

We have previously demonstrated that inhibition of ISR during the pre-diabetic phase (6-8 weeks of age) significantly reduces insulitis in NOD mice (8). In this context, we asked if macrophages act as key mediators in this ISR inhibition-driven reduction of immune infiltration. To determine the nature of immune cells in the insulitic regions of both vehicle and ISRIB-treated NOD mice, we performed Nanostring® spatial tissue proteomics using a pre-validated GeoMx® mouse immune cell panel. We did not observe any statistical differences in the immune cell subtypes within the insulitic regions following ISR inhibition (Figure 4A). However, in the islet area, as identified by positive staining for insulin, we observed an overall decrease in T cells (CD45+ cells) with a trend to reductions in CD3, CD4, and CD8 T cells (Figure 4B). Additionally, there was a trend towards a decrease in F4/80+ macrophages. Therefore, we next stained the pancreatic sections with F4/80 to directly measure the number of macrophages within the islet region. We observed a significant reduction in the number of macrophages after ISRIB treatment within the islet region (Figure 4C and D). Upon stratifying the islets into insulitic and non-insulitic categories, we observed a more pronounced reduction of macrophages within the insulitic islets, although a reduction was also evident in the non-insulitic islets (Figure 4E and F). The observed reduction in both insulitic and non-insulitic islets highlights the broader influence of ISR on macrophage dynamics within the pancreatic microenvironment.

### ISR Inhibition Increases PD-L1 Expression on Macrophages

Previous studies have shown that ISR inhibition in human islets enhances the expression of the immunosuppressive molecule PD-L1 on islet β cells (8, 9). PD-L1 is an immune checkpoint protein whose interaction with its receptor PD-1 on T cells reduces T cell cytotoxicity (19). One of the significant shared genes in the M1 transition in our RNA-sequencing was *Cd274* (encodes PD-L1), which increased more in the M1 ISRIB-treated group compared to the M1 vehicle group (Figure 2F). To build on these findings, we conducted immunofluorescence staining for PD-L1 on pancreatic sections from vehicle- and ISRIB-treated NOD mice. In this insulitic region where most macrophages are found, we observed an increase in PD-L1 levels after ISRIB treatment compared to treatment with vehicle (Figure 4G). Given the heterogeneity of cell types within these regions, we sought to determine if the increased PD-L1 expression was specific to macrophages. We analyzed PD-L1 expression in vehicle- and ISRIB-treated M1-polarized BMDMs in vitro and confirmed upregulation of PD-L1 by immunoblotting (Figure 4H). The mRNA levels of *Cd274* were not altered between the treatment groups (Figure 4I), suggesting a post-transcriptional mechanism. Our data suggests that ISR inhibition promotes PD-L1 expression in the remaining proinflammatory M1-like macrophages, potentially dampening T-cell activity.

## DISCUSSION

T1D is an autoimmune disease characterized by the destruction of insulin-producing β cells mediated by autoreactive T cells, with macrophages playing a pivotal role in initiating and amplifying this immune response (2). In this study, we sought to elucidate the role of the integrated stress response (ISR) in macrophages during T1D pathogenesis. We hypothesized that chronic activation of the ISR in macrophages promotes a proinflammatory state that contributes to the progression of insulitis and T1D. Our key findings are: (a) Inhibition of the ISR increases mRNA translation in proinflammatory macrophages; (b) ISR inhibition decreases macrophage migration, leading to fewer macrophages at sites of injury, including pancreatic islets; and (c) ISR inhibition increases PD-L1 expression on macrophages, potentially enhancing their ability to suppress T cell activity.

Macrophages are well-known drivers of early immune responses in T1D, infiltrating pancreatic islets before T and B cells and orchestrating the local inflammatory environment (20). Previous studies have shown that depletion of macrophages can prevent T1D onset in NOD mice, highlighting their central role in the disease’s progression (3, 4). Our findings build on these data by demonstrating that the ISR is a critical regulator of macrophage function. The ISR, through the phosphorylation of eIF2α, suppresses global mRNA translation while allowing selective synthesis of stress-related proteins (21). By inhibiting this pathway with ISRIB, we observed increased mRNA translation in M1-polarized proinflammatory macrophages, suggesting that the ISR serves as a regulatory checkpoint for protein synthesis during macrophage activation. This observation aligns with findings in other autoimmune conditions where heightened translational activity in immune cells is linked to inflammatory states (22).

The pancreatic lymph node (pLN) is required for the antigen presentation of APCs to T cells and removal of the pLN results in protection from diabetes in NOD mice (23). While it is still currently unknown how antigens flow from the pancreatic islet to the pLN, it is likely that resident islet APCs are migrating from the islet through the stroma and into the pLN to present the antigens to T cells (24). Our study shows that ISR inhibition disrupts this migratory capability. ISRIB-treated macrophages exhibited reduced expression of migration-related genes and impaired movement in both in vitro and in vivo models. This finding is consistent with reports that cellular stress pathways, including the ISR, influence cytoskeletal dynamics and cell motility (25). Notably, macrophage numbers within the pancreatic islets were significantly reduced following ISRIB treatment in NOD mice, particularly in insulitic islets, suggesting that ISR inhibition limits both macrophage recruitment and retention in inflamed tissues. This observation aligns with studies showing that targeting macrophage migration can reduce tissue-specific inflammation in autoimmune diseases (26), underscoring the potential of ISR modulation as a therapeutic strategy.

A particularly notable aspect of our findings is the increase in PD-L1 expression on macrophages following ISR inhibition. PD-L1 is an immune checkpoint molecule that plays a key role in maintaining self-tolerance by downregulating T cell activity (27). Previous work has demonstrated that blockade of PD-L1 accelerates diabetes in NOD mice, exacerbating the autoimmune response (28). Our observation that ISRIB increases PD-L1 levels on macrophages suggests that ISR inhibition not only alters macrophage function but may also enhance immune regulation through the suppression of autoreactive T cells. This finding situates the ISR as a potential modulator of immune checkpoint pathways, similar to its emerging role in cancer immunotherapy, where modulation of the ISR has been shown to influence antitumor immunity (29). The ability of ISRIB to increase PD-L1 expression offers a dual mechanism for therapeutic intervention—reducing macrophage-mediated inflammation while promoting T cell tolerance in the islets.

Our study, while providing new insights into the role of the ISR in macrophages during T1D, also highlights the need for further investigation. One limitation is our inability to distinguish between resident and infiltrating macrophages within the islets. The reduction in macrophage numbers observed in ISRIB-treated mice could reflect decreased migration of circulating monocytes/macrophages or changes in the resident macrophage population. Recent studies suggest that resident macrophages in pancreatic islets play distinct roles compared to infiltrating monocyte-derived macrophages, particularly in maintaining tissue homeostasis and regulating local immune responses (24). Future studies employing lineage tracing or single-cell RNA sequencing will be needed to clarify the contributions of these distinct macrophage populations to T1D progression.

In conclusion, our findings provide evidence that the ISR is a central regulator of macrophage function in the context of T1D. By modulating protein translation, macrophage migration, and immune checkpoint pathways, ISR inhibition may represent a therapeutic approach for limiting the proinflammatory actions of macrophages and promoting immune tolerance in the early stages of T1D. These insights not only advance our understanding of macrophage biology in autoimmune diabetes but also offer new avenues for targeting the ISR in inflammatory diseases more broadly.

## MATERIALS AND METHODS

### Animal Studies

All mouse experiments were conducted under specific pathogen-free conditions, with mice housed in a 12-hour light/dark cycle and given unrestricted access to food and water in accordance with protocols approved by the University of Chicago Institutional Animal Care and Use Committee. *C57BL/6J* (#664) and *NOD/ShiLtJ* (#1976) mice were obtained from Jackson Laboratories (Bar Harbor, ME). 6-week-old female *NOD/ShiLtJ* mice received intraperitoneal injections of either vehicle (5% DMSO, 2% Tween 80, 20% PEG400, and saline) or 2.5 mg/kg ISRIB (MedChemExpress; HY-12495) for two weeks (14). At the end of the study, the mice were euthanized, and pancreata from 4–5 mice per group were harvested, fixed in 4% paraformaldehyde, embedded in paraffin, and sectioned onto glass slides.

Zebrafish were maintained in 2 L tanks within a recirculating aquaculture system, held at a constant temperature of 28.5°C, and subjected to a 14-hour light/10-hour dark cycle in accordance with protocols approved by the University of Chicago Institutional Animal Care and Use Committee. The transgenic line *Tg*(*mpeg1:eGFP*)*^gl22^*(18) was originally obtained through the Zebrafish International Resource Center. Embryos were collected at spawning and incubated at 28.5°C in petri dishes containing egg water (0.1% Instant Ocean salts, 0.0075% calcium sulfate, and 0.1% methylene blue). At 24 hours post-fertilization (hpf), the egg water was replaced with fresh egg water supplemented with 0.003% 1-Phenyl-2-thiourea (PTU) to inhibit pigmentation.

### Macrophage Isolation

For in vitro studies using bone marrow-derived macrophages (BMDMs), femurs and tibias were excised and sterilized from 8-week-old *C57BL/6J* mice. BMDMs were isolated following a previously described method (30) and resuspended in RPMI supplemented with 10 mM HEPES, 100 U/mL penicillin/streptomycin, and 10% FBS (complete medium). Cells were plated and cultured in complete medium containing 30 ng/mL macrophage colony-stimulating factor (M-CSF; R&D Systems). The medium was refreshed every 3 days, and on day 6, the differentiated BMDMs were pretreated with either vehicle (DMSO) or 50 nM ISRIB for 1 hour, followed by treatment with 10 ng/mL LPS (Sigma) and 25 ng/mL IFN-γ (Prospec) for M1 polarization, 10 ng/mL IL-4 (R&D Systems) for M2 polarization, or control media for M0 (unpolarized) macrophages, in the continued presence of vehicle or ISRIB for 16 hours.

### RNA Isolation, Quantitative RT-PCR analysis, and RNA Sequencing

At the end of the polarization experiments, RNA was isolated using RNAeasy Mini kit (Qiagen) followed by cDNA synthesis using a High-Capacity cDNA Reverse Transcription kit (Applied Biosystems), according to manufacturer’s instructions. Quantitative PCR was conducted using SensiFAST™ SYBR**®** Lo-ROX Kit or Sensifast Probe Lo-ROX Kit (Thomas Scientific) on a CFX Opus system (Bio-Rad). The following mouse TaqMan gene probes were used: *Mif*: Mm01611157_g1; *Ccl5*: Mm01302427_m1; *Lgals3*: Mm00802901_m1; Cd274: Mm03048248_m1, and *Actb*: Mm01205647_m1 (Invitrogen). SYBR-green based quantitative PCR was performed for *Actb1* (*31*), *Nos2*, *Il6*, *Cxcr3* (*32*), *Ccr2* (*32*), *Cxcr1* (*33*), *Ccr4* (*32*), *Ccr7*, and *Slmaf9* (*34*) (primers details are listed in Supplemental Table S1). Relative gene expression was determined normalizing the average comparative threshold cycle (Ct) values of each gene to *Actb* expression. The normalized expression levels were then calculated relative to the averaged vehicle controls using the ΔΔCT method.

For RNA-sequencing, RNA was isolated from polarized and treated macrophages as described above. Samples were submitted to the University of Chicago sequencing core for library preparation and sequencing using the NovaSeq 6000® platform (Illumina). Raw sequencing data was analyzed using Rosalind (https://rosalind.bio/), leveraging the HyperScale architecture developed by ROSALIND, Inc. Reads were aligned to the Mus musculus genome (mm10 build) using STAR. Quantification of individual sample reads was performed with HTseq, and normalization was conducted via Relative Log Expression (RLE) using the DESeq2 R library. DESeq2 was also employed to compute fold changes, p-values, and optional covariate corrections. Pathway analysis was performed using Gene Ontology (GO). Previously published single-cell transcriptional data, available in GEO (GSE141786)(12), was reanalyzed using previously described methods using R (8). GSEA was performed using the msigdbr package in R to identify gene lists from specific gene sets, including the Hallmark Gene Set and GO Biological Processes.

### Protein Isolation, Immunoblotting, and SUnSET Assay

As described previously (8), whole-cell extracts were prepared using a lysis buffer (ThermoFisher) supplemented with HALT protease inhibitor cocktail (ThermoFisher). Proteins were then separated by electrophoresis using precast 4-20% tris-glycine polyacrylamide gels (Bio-Rad) and transferred to polyvinylidene difluoride (PVDF) membranes. Membranes were incubated overnight at 4°C with primary antibodies: anti-puromycin (Millipore; MABE343; 1:1000), anti-iNOS (Novus; NBPI-33780; 1:1000), anti-PD-L1 (abcam; ab213480; 1:1000), and anti-β-actin (Cell Signaling; 4970s, 3700s; 1:1000). Visualization and quantification were achieved using anti-rabbit or anti-mouse secondary antibodies (Li-Cor BioSciences; 1:10000), and blots were analyzed on the Li-Cor Odyssey system with data processed using the Odyssey Imaging software (Li-Cor Biosciences).

To determine the changes in protein translation, the SUnSET technique (13) was employed. Following treatment and polarization, BMDMs were incubated with 10 μg/mL puromycin for 15 minutes before protein isolation and puromycin intensity was normalized to β-actin levels for quantification.

### Flow Cytometry

Polarized and treated BMDMs were incubated on ice for 20 minutes with F4/80 (Biolegend; 123108) and CD11b (Biolegend; 101226) antibodies for surface marker staining. Following this, cells were permeabilized using Cytofix/Cytoperm (BD Pharmingen) and incubated on ice for an additional 20 minutes with CD206 (Biolegend; 141706) and iNOS (Invitrogen; 56-5920-82) antibodies for intracellular marker staining. All antibodies were used at a dilution of 1:100. Samples were analyzed using an Attune NxT Flow Cytometer (Thermo Fisher), and data were processed with FlowJo software (BD Biosciences). Antibody specificity was validated through flow cytometry controls, which included omitting either the primary or secondary antibodies, and employing gating strategies to eliminate background signals.

### Transwell Migration Experiments

The transwell migration assay was performed as described previously (5). Briefly, 2 × 10^5^ M1-polarized BMDMs were seeded into the upper chamber of a transwell apparatus equipped with an 8 μm pore size PET membrane (Corning), incubated for 4 h, and stained with crystal violet for visualization. 10% FBS served as a chemoattractant. Migrated cells were transferred to slides and imaged using a BZ-X810 fluorescence microscope (Keyence). Macrophage quantification was conducted through manual counting by blinded observers.

### Zebrafish Tailfin Injury Assay

Tailfin injury assays were performed as previously described (5). Briefly, at 3 dpf, zebrafish were treated with either DMSO control, 0.5, or 5 μM ISRIB for 18 h. Following treatment, zebrafish were immobilized with 0.01% tricaine and mechanical tailfin injuries were inflicted by excising the distal tips of the tailfins using a sharp scalpel. After injury, the embryos were transferred to fresh egg water in the presence of vehicle or different doses of ISRIB for 6 hours before fixation in 3% formaldehyde prepared in PEM buffer (100 mM PIPES, 1 mM MgSO4, 2 mM EGTA). The embryos were immunostained with TO-PRO-3 (Thermo Fisher; T3605; 1:1000) to identify the nuclei and images were captured using an A1 confocal microscope (Nikon). Image analysis was performed using ImageJ software. The number of macrophages that migrated to the area between the region distal to the notochord and the injury site was manually quantified by blinded observers.

### Immunostaining

For immunohistochemistry, paraffin-embedded pancreatic sections were stained with anti-F4/80 (Cell Signaling; D2S9R; 1:150) followed by the ImmPRESS*®* (Peroxidase) polymer anti-rabbit IgG reagent (Vector Laboratories). Visualization was achieved using the DAB peroxidase substrate kit (Vector Laboratories), and sections were counterstained with hematoxylin for cell nuclei (Sigma). Images were captured using a BZ-X810 fluorescence microscope (Keyence). For analysis, three sections per mouse, spaced 100 μm apart, were examined and averaged. Macrophage quantification was performed manually by blinded observers.

For immunofluorescence staining, pancreatic tissues were stained with anti-PD-L1 (R&D Systems; AF1019; 1:200) and anti-insulin (Dako; IR002; 1:4) antibodies. Highly cross-adsorbed Alexa Fluor secondary antibodies (ThermoFisher; 1:500) were used for detection, and nuclei were stained with DAPI (ThermoFisher). Images were acquired using an A1 confocal microscope (Nikon). Background subtraction was performed using ImageJ software. The specificities of the antibodies were confirmed in tissue sections in which the primary or secondary antibody were omitted and no relevant signal was observed.

### NanoString® Spatial Proteomics

Paraffin-embedded pancreatic tissues were analyzed using Nanostring*®* spatial proteomics as described previously (8, 9). Briefly, morphological staining was carried out with AF-647 conjugated insulin (Cell Signaling; 9008s; 1:400) and SYTO13 for nuclear visualization. Spatial hybridization utilized a pre-validated mouse GeoMx Immune Cell Panel (NanoString; GMX-PROCONCT-MICP). Background correction was achieved using IgG antibodies (Rb IgG, Rat IgG2a, Rat IgG2b). Regions of interest (ROI) were selected based on morphology, specifically focusing on 5-6 islets exhibiting insulitis per mouse. These ROIs were then further divided into insulitic and insulin-positive areas within each islet. Oligonucleotides from these segmented regions were photocleaved and quantified using the nCounter system (NanoString). Data analysis was performed using NanoString*®* software, incorporating scaling adjustments for variations in tissue surface area and depth. After scaling, read counts were normalized to housekeeping genes, and background signals were subtracted using IgG markers.

### Statistical Analyses

All data are reported as mean ± SEM. To compare multiple groups, either one-way ANOVA or repeated measures ANOVA was conducted, followed by Tukey’s post-hoc test. For two-group comparisons, two-tailed unpaired or paired Student’s t-tests were utilized. Statistical analyses and graphical representations were generated using GraphPad Prism v10. Statistical significance was established at *P* < 0.05.

## SUPPORTING INFORMATION

This article contains supporting information.

## ACKNOWLEDGEMENTS

We thank Jennifer Nelson, Sveltana Navitskaya, Sarida Pratuangtham, Advaita Chakraborty, and Kayla Figatner for their technical help.

This work was supported in part by National Institutes of Health grants R01 DK135832 (to SAT), U01 DK127786 and R01 DK060581 (both to RGM), Quad Summer Scholarship award (to JEW), Breakthrough T1D postdoctoral fellowship (3-PDF-2023-1326-A-N) and Diabetes Research Connection awards (both to CM). This study utilized Diabetes Center core resources supported by National Institutes of Health grant P30 DK020595 (to the University of Chicago) and utilized services of the University of Chicago Histology and Genomics Cores.

## AUTHOR CONTRIBUTIONS

Jiayi E. Wang: Methodology, Investigation, Funding Acquisition, Visualization, Writing – original draft

Charanya Muralidharan: Conceptualization, Methodology, Investigation, Funding Acquisition, Visualization, Writing – original draft

Titli Nargis: Investigation, Writing – review and editing

Jacob R. Enriquez: Investigation, Writing – review and editing

Ryan M. Anderson: Supervision, Writing – review and editing

Raghavendra G. Mirmira: Conceptualization, Supervision, Project Administration, Funding Acquisition, Writing – review and editing

Sarah A. Tersey: Conceptualization, Supervision, Project Administration, Funding Acquisition, Visualization, Writing – original draft

## DATA AVAILABILITY

RNA sequencing data have been uploaded to the public repository GEO (GSE278047). Any additional information required to reanalyze the data reported in this paper is available from the corresponding author upon request.

