## Supporting Information for "The integrated stress response promotes macrophage inflammation and migration in autoimmune diabetes"

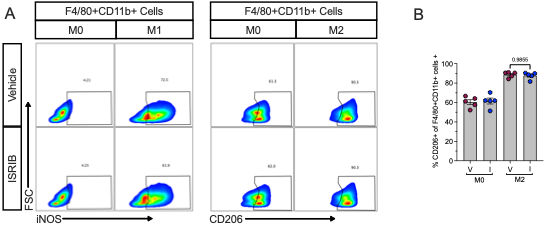

**Supplemental Figure S1: ISR Inhibition Selectively Modulates Propensity for Classical Polarization.** (**A**) Representative contour plots gating of iNOS+ F4/80+ CD11b+ cells (*left panels*) and representative contour plots gating of CD206+ F4/80+ CD11b+ cells (*right panels*). (**B**) Quantification of CD206+ (M2-polarization marker) cells as a percentage of F4/80+ CD11b+ cells; *n* = 5 biological replicates. Statistical test: one-way ANOVA. Data are represented as mean ± SEM.

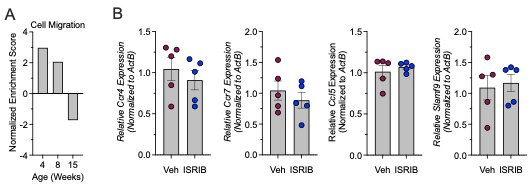

**Supplemental Figure S2: Temporal Dynamics of Macrophage Migration and Migration Genes Unaltered by ISR Inhibition.** (**A**) GSEA of the macrophage cell population of the scRNA-Seq dataset identified in **Fig.** **1C** for cell migration. (**B**) Relative mRNA levels of macrophage migration related genes measured by qRT-PCR normalized to *Actb* in M1-polarized BMDMs from male 8-week-old *C57Bl/6J* mice; *n* = 5 biological replicates. Statistical tests: Paired t-tests. Data are represented as mean ± SEM.

**Table S1: RT-PCR Primer Sequences**

| Gene | Forward | Reverse |
| --- | --- | --- |
| *Actb* | 5'-GGCTGTATTCCCCTCCATCG-3' | 5'-CCAGTTGGTAACAATGCCATGT-3' |
| *Nos2* | 5'-GAGACAGGGAAGTCTGAAGCAC-3' | 5'-CCAGCAGTAGTTGCTCCTCTTC-3' |
| *Il6* | 5'-TACCACTTCACAAGTCGGAGGC-3' | 5'-CTGCAAGTGCATCATCGTTGTTC-3' |
| *Cxcr3* | 5’-GCTGCTGTCCAGTGGGTTTT-3’ | 5’-AGTTGATGTTGAACAAGGCGC-3’ |
| *Ccr2* | 5’-AACAGTGCCCAGTTTTCTATAGG-3’ | 5’-CGAGACCTCTTGCTCCCCA-3’ |
| *Cxcr1* | 5’-CCGTCATGGATGTCTACGTG-3’ | 5’-CAGCAGCAGGATACCACTGA-3’ |
| *Ccr4* | 5’-AAGAAGAACAGAGCAGTGCGC-3’ | 5’-GCGACCAGAAGCCGAGG-3’ |
| *Ccr7* | 5’-CGGGATGCTGCTGCTCCTATGC-3’ | 5’-TCCTCGCCGCTGTTCTTCTGGA-3’ |
| *Slamf9* | 5’-CCTAGCCAGCTGACCAAGTC-3’ | 5’-GCGTTGTAAAGCCCTGAGTC-3’ |
